# Environment-dependence of the expression of mutational load and species’ range limits

**DOI:** 10.1101/2021.09.15.460519

**Authors:** Antoine Perrier, Darío Sánchez-Castro, Yvonne Willi

**Affiliations:** Department of Biology, University of Virginia, 485 McCormick Road, Charlottesville, VA 22904, USA; Department of Environmental Sciences, University of Basel, Schönbeinstrasse 6, 4056 Basel, Switzerland

**Keywords:** *Arabidopsis lyrata*, environmental stress, genetic drift, range limit, heterosis, mutational load

## Abstract

Theoretical and empirical research on the causes of species’ range limits suggests the contribution of several intrinsic and extrinsic factors, with potentially complex interactions among them. An intrinsic factor proposed by recent theory is mutational load increasing towards range edges because of genetic drift. Furthermore, environmental quality may decline toward range edges and enhance the expression of load. Here we tested whether the expression of mutational load associated with range limits in the North American plant *Arabidopsis lyrata* is enhanced under stressful environmental conditions by comparing the performance of within- *versus* between-population crosses at common garden sites across the species’ distribution and beyond. Heterosis, reflecting the expression of load, increased with heightened estimates of genomic load and with environmental stress caused by warming, but the interaction was not significant. We conclude that range-edge populations suffer from a twofold genetic Allee effect caused by increased mutational load and stress-dependent load linked to general heterozygote deficiency, but no synergistic effect between them.

## Introduction

The geographic distribution of species can be constrained by several factors, and they may interact in complex manners (Gaston, 2009; Roy et al., 2009; Sexton et al., 2009; Connallon & Sgrò, 2018; Willi & Van Buskirk, 2019). An important factor put forward by relatively recent theory is the accumulation of mutational load in range-edge populations due to a history of enhanced genetic drift (Peischl et al., 2013; Henry et al., 2015; Willi, 2019). In parallel, environmental conditions may degrade toward the range limits of a species (Brown, 1984; Cahill et al., 2014; Hargreaves et al., 2014; Lee-Yaw et al., 2016). In addition to their respective negative effect on population performance, mutational load and environmental stress may interact synergistically in their effect on performance, as was suggested and found for inbreeding depression (Roff, 1997, pages 285-338; Reed et al., 2012). The contribution of mutational load to range limits could therefore be much more substantial than expected by previous theoretical and empirical work and relevant for shaping the distribution of many species.

Two types of demographic scenarios at range edges are predicted to lead to increased mutational load. First, theoretical models predict that range expansion through serial demographic bottlenecks enhances genetic drift and lowers the efficacy of purifying selection, which leads to the accumulation of deleterious mutations (Peischl et al., 2013; 2015; Peischl & Excoffier, 2015). High load at the expansion front is predicted to slow down the spread of a species or halt it if genome-wide recombination is low (Peischl et al., 2015). The so-called expansion load is predicted to involve mainly recessive deleterious mutations (Peischl & Excoffier, 2015), to establish even in the absence of environmental gradients, and to persist in populations for thousands of generations (Peischl et al., 2013). Second, if populations at range edges are small and isolated, genetic drift is also enhanced and increases mutational load. Simulations showed that when the carrying capacity of suitable habitat declines along a line of habitat patches, the occupied range becomes shorter than the distribution of habitat would suggest because of mutation accumulation (Henry et al., 2015). The effect of load is larger if dispersal and population growth rates are low (Henry et al., 2015), and this scenario of mutational load limiting ranges may apply best to rear edges of species’ distribution (Hampe & Petit, 2005). Genomic evidence for the accumulation of mutational load under range expansion was found for humans (Lohmueller et al., 2008; Peischl et al., 2018) and under expansion and rear-edge isolation for plants (Zhang et al., 2016; González-Martínez et al., 2017; Willi et al., 2018; Koski et al., 2019). At range limits, the environment may also be more stressful. Empirical studies document a general decline in habitat suitability toward range limits in many taxa (Sexton et al., 2009; Pironon et al., 2017). A rather consistent and pronounced effect of declining habitat marginality was found in species that were experimentally transplanted beyond their range edges as their lifetime performance generally declined compared to sites within the range (Hargreaves et al., 2014). The decline in performance was found to coincide with a decline in predicted habitat suitability revealed by niche modelling (Lee-Yaw et al., 2016). The two meta-analyses suggest that the decay in habitat suitability is rather general. Such unfavourable conditions may act in isolation and lower population performance, or they may enhance the expression of mutational load at range edges, further reducing population performance.

Potential mechanisms for a synergistic interaction between mutational load and environmental stress have rarely been explored *per se*, but have been mainly discussed and tested in the context of inbreeding depression (Armbruster & Reed, 2005; Willi et al., 2007a; Fox & Reed, 2010). Three mechanisms have been proposed (Reed et al., 2012). The first is that stress induces the expression of deleterious mutations that are silent under benign conditions (Kondrashov & Houle, 1994; Elena & de Visser, 2003). For example, in *Drosophila*, recessive alleles were linked to increased mortality in inbred lines under temperature stress (Vermeulen & Bijlsma, 2004). Inbreeding and extreme stress impede the function of heat shock proteins such as Hsp90, which are essential in buffering the expression of deleterious mutations (Rutherford & Lindquist, 1998; Queitsch et al., 2002; Bergman & Siegal, 2003). A second mechanism is that mutational load may lower stress resistance or tolerance, by destabilizing cellular homeostasis under stress, or affecting tissue and genome repair after stress exposure (Agrawal & Whitlock, 2010; Reed et al., 2012). For example, hybrids of *Arabidopsis thaliana* expressed more metabolites from central pathways linked to higher freezing tolerance or disease resistance compared to inbred lines (Korn et al., 2010; Yang et al., 2015), and the regulation of stress-response pathways led to higher recovery after stress (Miller et al., 2015). The third mechanism is that stress does not increase the expression of load *per se*, but rather increases phenotypic variance and with it the opportunity for relative fitness measures to differ more between inbred and outbred individuals (Waller et al., 2008). Reed et al. (2012) found that variation in inbreeding depression was mainly predicted by stress itself and only to a weaker extent by increased phenotypic variance.

In this study, we tested for stress dependence of the expression of mutational load at the range limits of North American *Arabidopsis lyrata* subsp. *lyrata* (L.). The species has been strongly impacted by the last glacial maximum (LGM), with more than half of its contemporary distribution resulting from rapid post-glacial range expansion from two separate refugia (Griffin & Willi, 2014; Willi et al., 2018). Both range expansion and rear-edge dynamics were found to be associated with an increase in presumably deleterious mutations assessed on the level of DNA sequences, leading to a current pattern of heightened deleterious mutations in range edge populations (Willi et al., 2018). Another study showed that this heightened genomic load was correlated with expressed load, estimated by heterosis – the fitness increase of between-compared to within-population crosses (Perrier et al., 2020). Here we tested whether the expression of mutational load, again estimated by heterosis, additionally depended on the interaction between genomic load and environmental stressfulness in the direction of a positive synergistic effect. Offspring of within-(WPC) and between-population crosses (BPC) were raised in a latitudinal transplant experiment with five common garden sites within and beyond the distribution range (plant data as in Perrier et al. (2020)). Environmental stressfulness was estimated by (a) the differences in climatic conditions experienced by populations in each common garden relative to their site of origin, assuming local adaptation to climate (Hoffmann & Hercus, 2000), and by (b) the relative performance of each population at a garden site compared to the best garden. Finally, we tested whether heterosis in range-edge populations – expressing the highest mutational load compared to populations from elsewhere of the range – was enhanced when offspring of crosses were transplanted to environments beyond range limits.

## Materials and Methods

### Experimental design

#### Populations

Twenty populations of *A. lyrata* subsp. *lyrata* from across the species’ distribution were selected for producing WPC and BPC (Fig. 1; Table S1). The populations represented the two main ancestral clusters (Griffin & Willi, 2014; Willi et al., 2018), the latitudinal range, and variation in the genomic signatures of mutational load (Willi et al., 2018). Our sampling also represented the two mating systems: predominant outcrossing and predominant selfing (MO2, ON1, ON11). In the study system, selfing populations are mainly restricted to the edges of distribution of the two ancestral clusters (Griffin & Willi, 2014). They have an especially high genomic load, on the one hand, because of their expansion history, and on the other hand, because of the mating system shift (Willi et al., 2018; Perrier et al., 2020). Replicate selfing populations were included to increase the range of genomic load found in the study system. Eighteen populations (nine per cluster) were considered target populations in the assessment of expressed mutational load, and two were partner populations for BPC (NY1, eastern cluster, and IA1, western cluster). Partner populations were selected because of their core location in each genetic cluster, high genomic diversity, and history of little genetic drift. Between-population crosses with these partner populations were therefore expected to overcome mainly the load of target populations. Seeds of individual plants had been collected in each population over an area of about 450 m^2^.

**Figure 1:**
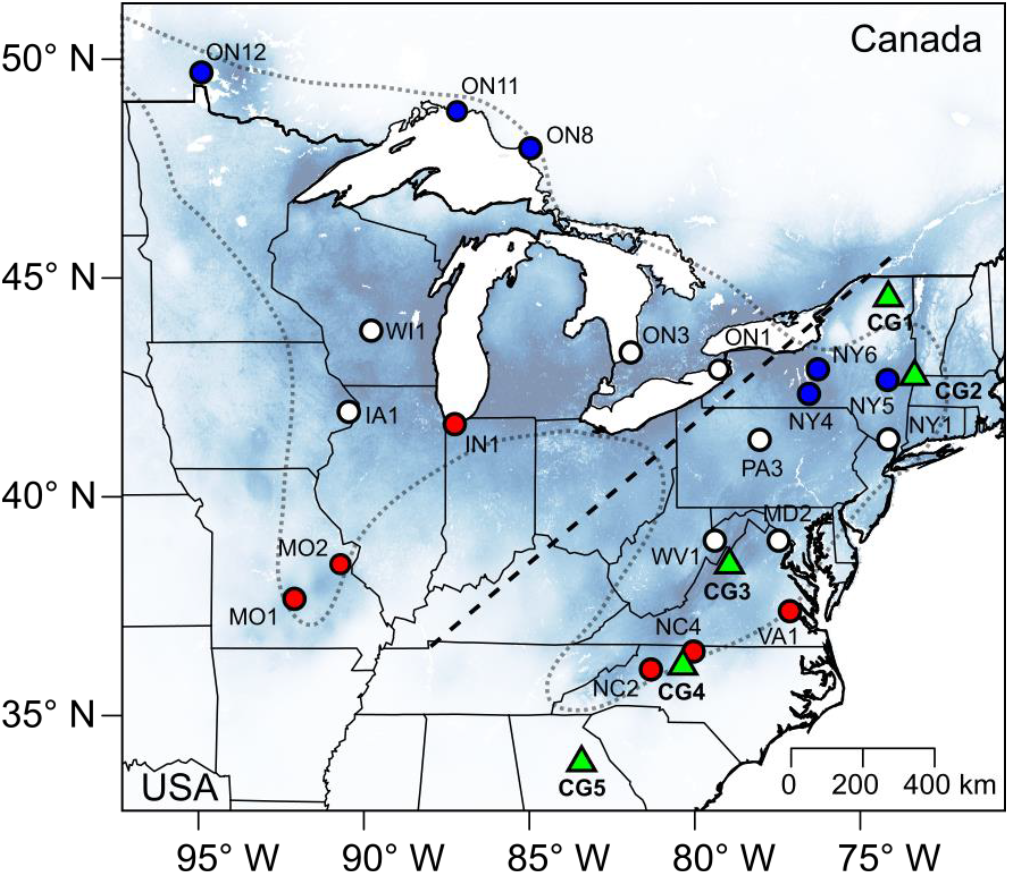
Distribution of *Arabidopsis lyrata* in eastern North America, with information on habitat suitability, the location of the 20 populations studied, and the 5 common garden sites. The range of *A. lyrata* is represented by the dotted line, habitat suitability by shades of blue, with darker blue indicating higher suitability (Lee-Yaw et al., 2018). Populations are shown by circles, with abbreviations for state (USA) or province (Canada) and a number (Willi et al., 2018). Blue and red circles represent northern- and southern-edge populations in our analysis. Green triangles represent the five common garden (CG) sites; numbers added to labels are in sequence of north to south. State outlines for the USA are shown, and the split between eastern and western genetic cluster is represented by the dashed line. Of the 20 populations, two were used as partner-populations for between-population crosses, NY1 for crosses with eastern populations, and IA1 for crosses with western populations.

#### Crossing

One plant of each of 26 randomly selected and presumably unrelated seed families per population were raised under controlled, indoor conditions. Plants were randomly appointed to “mothers”/pollen recipients (12), “fathers”/pollen donors (12), and backups (2). Each “mother” was crossed using pollen from one “father” of the same population (WPC). For target populations, each “mother” was additionally crossed using pollen from one “father” of the partner population (BPC).

#### Crosses were non-reciprocal

Pollination was performed at the bud stage to prevent unintended cross- or self-pollination. We repeated each cross-combination to obtain enough healthy-looking seeds to perform the transplant experiment (min. 60). In the case of systematic cross failure (overall fractions of failed crosses were similar between WPC and BPC, about 8%), we replaced the father plant with a backup plant. In total, 401 family-cross type combinations were used for the transplant experiment (Table S2).

#### Transplant experiment

Five common garden sites (CG) were selected along a *ca.* 1400 km latitudinal gradient crossing the distribution of *A. lyrata* in the eastern USA (Fig. 1; Table S3 with WorldClim data v 2.0 (Fick & Hijmans, 2017)). The five sites represented the centre of the species’ range (CG3, Harrisonburg, VA), the southern and northern range edges (CG4, Winston-Salem, NC, and CG2, Williamstown, MA, respectively), and areas beyond the edges (CG5, Athens, GA, and CG1, in the Adirondacks, NY). In 2017, in each common garden, two seeds (one if few seeds were available) of each cross were sown in pots (5.7 cm wide and 5.9 cm deep, with holes at the bottom) randomly distributed within each of three spatial blocks, resulting in three to six seeds per cross in each common garden (more details in Perrier et al. (2020)). A total of 12,933 seeds were sown across the five common gardens. Five data loggers (iButton®, Maxim Integrated Products, Inc) monitored air temperature every hour (1.5 m above ground, in the shadow) in each site for the duration of the experiment.

#### Performance

Germination was recorded for each seed three days a week until peak germination was over (four to five weeks) and afterward once a week until thinning, starting 11 weeks after sowing. Seedlings were randomly thinned to one per pot. Reproductive output was assessed several weeks after peak flowering in 2018 and 2019 and calculated for each individual as the sum of fruits, pedicels (flowers that did not develop into fruits), flowers, and flower buds. Flowers that did not develop into fruits were included because they contributed to the performance capacity of plants, and their pollen may have fertilized ovules on other plants. However, seeds per fruit were not estimated as pollen transfer was not controlled for in the experiment. In an additional experiment, we assessed seed survival over winter, with seeds from the same crosses as used in the transplant experiment, to estimate how seeds that had not germinated the first fall could have contributed to population performance. For each population-cross type combination, we pooled 100 seeds from five to twelve seed families, split them in 10 bags (made of nonwoven polypropylene-felt, 40 g/m2 to allow air and moisture exchange) and placed 2 bags per garden on bare ground next to pots in October 2018. Bags were retrieved in late spring 2019 and visually screened for germinated seedlings and seeds. Remaining seeds were tested for germination in the laboratory. Seed survival over the winter was calculated for each bag as the fraction of germinated seedlings outdoors and in the lab and used for estimating population growth rate (for each combination of population, cross type, and garden).

### Statistical analysis

Initial analyses were based on *Multiplicative performance* (*MP*) calculated on the level of the pot as the product of the fraction of germinated seeds and the sum of total reproductive output recorded until year 3 (year 2 for CG 3 that had to be given up early). We tested for a general effect of cross type (BPC compared to WPC, which were coded as 1 and 0, respectively), genomic load, environmental stress, and all two- and three-way interactions between them. Genomic load was an estimate of mutational load based on nuclear DNA sequences. Population pools of equimolar DNA of 25 plants per population were whole-genome sequenced, nuclear single-nucleotide polymorphisms were retrieved, and those in coding regions were split into non-synonymous and synonymous. For each population, the ratio of non-synonymous polymorphic sites to synonymous polymorphic sites adjusted for their mean derived allele frequency (relative to *A. thaliana*), *P*_n_*f*_n_/*P*_s_*f*_s_ was calculated (Willi et al., 2018). This estimate of mutational load should depict the excess number and frequency of derived and presumably deleterious mutations due to genetic drift. The estimate of mutational load correlated well with population genomic diversity in intergenic DNA, which itself is affected little by interpopulation gene flow in the species. Between-population crosses were assigned the genomic load of the mother population as all BPC shared the same pollen donor population within a genetic cluster.

Environmental stress was depicted by the difference in climate between garden site and site of origin of a population (Hoffmann & Hercus, 2000, Rutter and Fenster, 2007). Distribution modelling had shown that average minimum temperature in March and April described best the niche and latitudinal range limits of *A. lyrata* (Lee-Yaw et al., 2018). Therefore, environmental stress was calculated as the difference in average minimum temperature in early spring (March and April) between common garden sites and site of origin of populations: Δ *T*_min_ *= T*_min CG_ - *T*_min origin_. *T*_min CG_ was calculated based on records of temperature loggers (Table S4), and *T*_min origin_ based on the WorldClim 2.0 database (spatial resolutions of 30 seconds (Fick & Hijmans, 2017); Table S1) and averaged over target and partner population for BPC. Positive and negative values of Δ *T*_min_ indicated transplanting toward warmer and colder temperatures, respectively. We assumed that both positive and negative deviations could be perceived as stressful, resulting in a bell-shaped relationship between Δ *T*_min_ and performance. Therefore, we included the square term of Δ *T*_min_ in the model. Evidence for a positive synergistic effect between the expression of genomic load – the superior performance of between- to within-population crosses – and environmental stress was assumed to be given by the positive three-way interaction coefficient. Analyses were performed in a Bayesian framework on 10 parallel chains (no. of iterations: 50000, burnin: 5000, thinning: 100, leading to 450 posterior estimates; code in Method S1), with the package *MCMCglmm* (Hadfield, 2010; Hadfield, 2019) in *R* (R Core Team, 2021). The model accounted for the zero-inflation of *MP* by considering a Gaussian process (*MP* > 0, hereafter referred to as the log-normal process) and a logistic process (probability of 1 assigned to *MP* > 0). Random effects were mother plant (that was crossed) nested within maternal population, and maternal population, for which variance in intercepts and slopes of cross type were estimated; and block nested within common garden and common garden. Covariates were mean-centred, and square terms were calculated based on mean-centred values.

Further analyses were – for ease of interpretation – performed on heterosis estimated on the level of the population to determine the best model on the expression of load under environmental stress. Heterosis was calculated as the increase in performance of BPC relative to WPC: (*W_BPC_ - W_WPC_)/W_WPC_*, where *W_WPC_* and *W_BPC_* were population averages based on seed family averages in a common garden. *W_WPC_* was further averaged across target and partner populations. Heterosis was also assessed for the finite rate of increase per year, *λ*, from stage-classified matrices (Caswell, 2001) based on the mean performance of plants of each population-cross type-garden group, over the three years (detailed in Perrier et al. (2020)). These matrices comprised the stages: (1) healthy seeds in year 1, (2) reproducing individuals of year 2, (3) reproducing individuals of year 3, with a one-year projection interval between each stage. Heterosis estimates were log_10_-transformed and analysed by hierarchical mixed-effects models based on restricted maximum likelihood using the packages *lme4* (Bates et al., 2015) and *LmerTest* (Kuznetsova et al., 2017) in *R*. Fixed effects were the genomic estimate of mutational load of the mother population, environmental stress, and interactions between them. Crossed random effects were population and common garden (detailed in Method S2A). Here, Δ *T*_min_ was calculated with *T*_min origin_ again averaged over target and partner population.

To explore the effect of stress, we depicted it by either signed or absolute Δ *T*_min_, included an orthogonal quadratic term or not, and included interaction terms or not. We performed model selection based on the corrected Akaike information criterion, AICc (Sugiura, 1978). We selected the model with the lowest AICc value, with the additional criterion that its AICc had to be at least 2 units below the next better model. For the best model, we tested the significance of each fixed effect with a likelihood-ratio *χ*^2^ test using the *Anova* function (type III) of the package *car* (Fox & Weisberg, 2019). We also used the same model to test for effects on *W_WPC_* to estimate the likely effect genomic load may have under stress in the natural populations. A confounding role of genetic incompatibility estimated by neutral genetic divergence and of divergent adaptation between target and partner populations estimated by environmental divergence could be excluded. Details on these analyses are presented in Appendix S1. We repeated the analysis on heterosis on a second estimate of stress, the relative decline in population-level performance at a site relative to the best site (1 - *W*_WPC_/*W*_max_) (Reed et al., 2012).

Stress varied between 0 (conditions leading to maximal performance) and 1 (conditions with *W*_WPC_ = 0). Relative performance was then averaged over both target and partner populations. Finally, we tested whether heterosis was higher when range-edge populations (with highest load) were transplanted beyond the range limits compared to sites at the range limits. We assessed this effect in two distinct datasets: one considering the six northern edge populations (ON8, ON11, ON12, NY4, NY5, and NY6; in blue in Fig. 1) raised at the northern edge site (CG2) and beyond the northern edge (CG1), and one considering the six southern edge populations (MO1, MO2, IN1, NC2, NC4, and VA1; in red in Fig. 1) raised at the southern edge site (CG4) and beyond the southern edge (CG5). For both datasets, log_10_-transformed heterosis was tested in hierarchical mixed-effect models based on restricted maximum likelihood. Transplant was a fixed effect (edge = 0), and population was a random effect (detailed in Method S2B).

## Results

Analyses on the level of the pot revealed a significant positive effect of cross type on performance in the logistic process (probability of 1 assigned to *MP* > 0, Table 1). Performance was higher in between-compared to within-population crosses (BPC and WPC), supporting a general heterosis effect. Performance was also significantly negatively affected by genomic load and by a positive cross type-by-genomic load interaction in the log-normal process (*MP* > 0). Performance generally declined with increasing genomic load but less in BPC than WPC. Performance was further marginally negatively affected by the linear term of stress (Δ *T*_min_) and significantly negatively affected by the quadratic term of stress (Δ *T*_min_^2^), both in the log-normal and logistic process. Performance generally declined when conditions in the common garden diverged from conditions at the site of origin (Δ *T*_min_ ≠ 0). Furthermore, the interaction cross type-by-Δ *T*_min_ was significantly positive for the logistic process, indicating that the decline due to stress was stronger for WPC compared to BPC, especially under warmer conditions (Δ *T*_min_ > 0). The logistic process determining performance was also significantly affected by a negative interaction between genomic load and Δ *T*_min_. Performance of populations with higher load declined more when exposed to warmer temperatures but less when exposed to colder temperatures. Finally, three-way interactions between fixed effects were not significant.

**Table 1:**
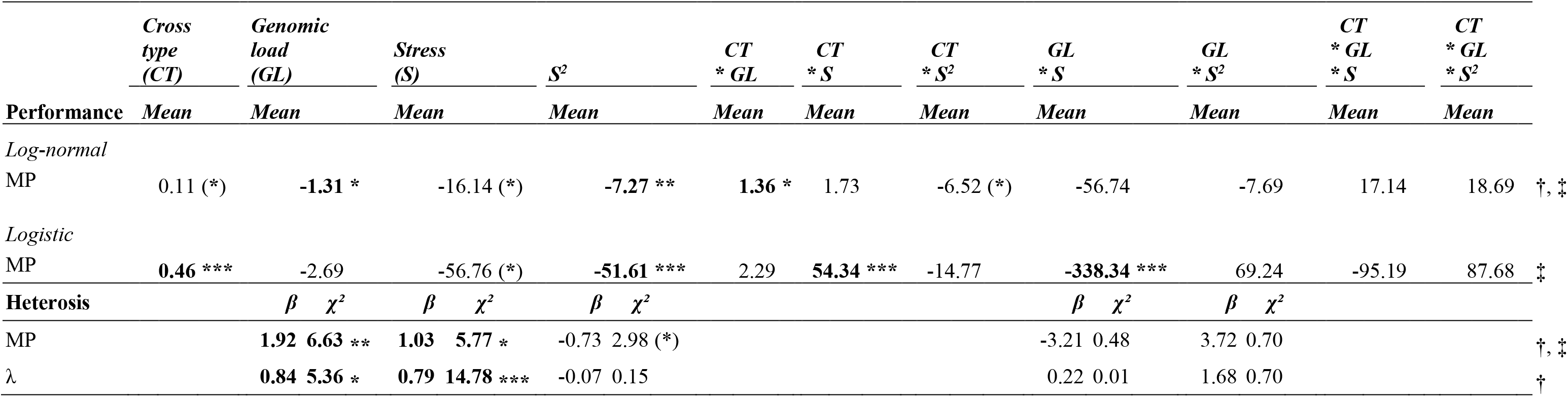
Results of model testing for the effect of cross type (between-compared to within-population crosses), genomic estimate of mutational load, environmental stress (Δ *T*_min_ *= T*_min CG_ - *T*_min origin_) and their interactions on *multiplicative performance* (*MP*), or on heterosis based on *MP* and the finite rate of increase (*λ*), at the five common garden sites In the models testing multiplicative performance (MP, log_10_-transformed if > 0), MP was assumed to follow a Gaussian distribution with 0-inflation. Therefore, the model assessed all fixed and random effects for their importance in both the logistic process (binary variable depicting germination combined with survival and the capacity to initiate flowering) and the Gaussian process (total number of flowers during one or two reproductive seasons). For this model, estimates of coefficients are modes of MCMC samples from the posterior distribution of parameters (*mean*). The logistic part of the model predicts non-zeros in the distribution on the logit scale. In the model testing population heterosis, estimates of population heterosis were log_10_-transformed prior to analysis. Stress, *Δ T*_min_, was an average over target and partner population. Each model testing heterosis was optimized with the *bobyqa* optimizer to improve convergence. Test statistics include regression coefficient (*β*) and the *χ*²-value of each fixed effect. For heterosis based on MP and on ***λ***, the marginal *R*^2^ of the model was of 0.18 and 0.23 respectively, and the conditional *R*^2^ of the model was of 0.37 and 0.46 respectively. Genomic load and stress were standardized prior to analyses (mean = 0). Model fits with significant (positive) intercept are indicated by †. Estimates and regression coefficients with *P*-values < 0.05 are written in bold; significance is indicated: (*) *P* < 0.1, * *P* < 0.05, ** *P* < 0.01, *** *P* < 0.001. Results for random effects are not shown. For one of the five common gardens (CG3), the experiment stopped early and variables consider performance to year 2 only (indicated by ‡).

Results on population-level heterosis based on *MP* or *λ* revealed significant positive associations between genomic load and heterosis and between the linear term of stress and heterosis (Table 1, Fig. 2a), in line with the significant cross type-by-genomic load and cross type-by-Δ *T*_min_ interactions in the individual-level analysis. Heterosis was highest in populations with more mutational load, and in populations raised under warmer conditions than at their site of origin (Δ *T*_min_ > 0). Neither the quadratic term of stress nor the interactions between stress and genomic load were significant. Heterosis based on *MP* and *λ* in the five common gardens ranged from −0.96 to 23.50 (mean: 1.88) and −0.53 to 7.29 (mean: 0.73) respectively (Table S5).

**Figure 2:**
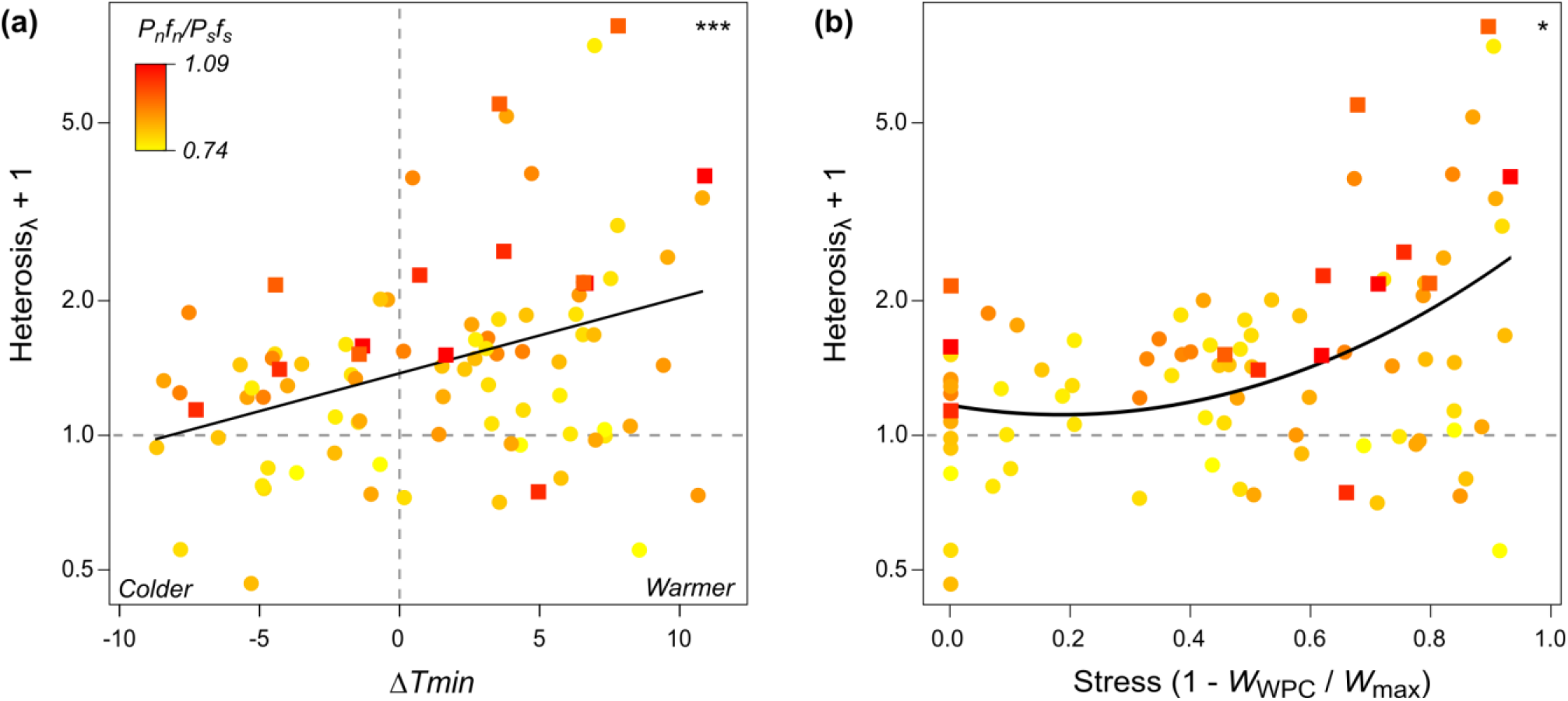
Stress dependence of the expression of mutational load estimated by heterosis. Population heterosis was estimated based on the finite rate of increase, *λ*, of within-population crosses and between-population crosses with a partner population. Outcrossing populations are indicated by dots, selfing populations by squares. Stress was depicted as either the difference in minimum temperature in early spring between common garden and the site of origin of a population (**Δ** *T*_min_ = *T*_min CG_ - *T*_min origin_; positive values to the right of the vertical dashed line indicate a warmer environment) **(a)** or the relative decline in population-level performance *W* in a garden relative to the garden of highest performance for that population (1 - *W*_WPC_/*W*_max_) **(b)**. For both stress estimates, means of target and partner populations were taken. The horizontal dashed lines indicate when heterosis drops below 0, and outbreeding depression dominates. The black lines represent model-predicted relationships between heterosis and environmental stress (test statistics in Tables 1, Table S10; * *P* < 0.05, *** *P* < 0.001). Genomic mutational load, the ratio of genome-wide non-synonymous to synonymous polymorphic sites adjusted by their mean derived frequency, *P*_n_*f*_n_/*P*_s_*f*_s_ (Willi et al., 2018), is represented in shades of yellow (low) to red (high).

Model selection pointed to this full model on heterosis being the best supported (Table S6). Among the models compared, the one with genomic load, the signed term of stress (Δ *T*_min_), its square term, and interactions between genomic load and stress had by far the lowest AICc (next better model with + 6.7 AICc units). Similar to the analysis of heterosis, its component of *W_WPC_* revealed an effect of genomic load, Δ *T*_min,_ and Δ *T*_min_^2^ (Table S7), but no effect of the genomic load-by-Δ *T*_min_ interaction. The performance of WPC declined with increasing genomic load and, in an accelerating manner, with warmer conditions compared to the site of origin of populations (Fig. S1). The model-predicted maximal decline in WPC performance for a population with average genomic load was 92% when exposed to warmer temperatures (baseline of Δ *T*_min_ = 0; Table S8).

Analyses on the effect of stress expressed by performance decline on heterosis revealed similar results as those based on Δ *T*_min_. The best model predicting heterosis included the linear and quadratic terms for stress and interactions between stress and genomic load (next better model with + 7.3 AICc units; Table S9). Heterosis was significantly positively affected by genomic load (marginal for *λ*) and both the linear (except *MP*) and the quadratic term of stress (Table S10) but was again unaffected by interactions. Heterosis based on *MP* increased almost symmetrically toward low (*W*_WPC_ *= W*_max_) and high stressfulness (*W*_WPC_ *= 0*; Fig. S2), while heterosis based on *λ* increased more strongly toward high stressfulness (Fig. 2b). Finally, heterosis of range edge populations did not significantly differ between the garden sites at the range edge and the garden sites beyond the range, neither for the southern nor for the northern range edge (Table S11, Fig. S3).

## Discussion

Recent evolutionary theory proposes that species’ range limits may result from high mutational load in range-edge populations due to prolonged exposure to genetic drift caused by past range expansion or rear-edge demographics (Willi, 2019). Here we tested whether the expression of such load is enhanced by environmental stress in a large-scale transplant experiment with *Arabidopsis lyrata*. To express load, we used heterosis based on the multiplicative performance of between-compared to within-population crosses, assuming that load was mainly due to recessive deleterious mutations. To estimate stress, we used divergent thermal conditions during a critical part of the year between gardens and sites of origin of populations or relative performance decline of each population. Expressed mutational load was stronger in populations with higher deleterious counts inferred from coding DNA sequences, as previously found (Willi et al., 2018; Perrier et al., 2020). However, stress did not clearly synergistically interact with genomic load to enhance heterosis. Nonetheless, we found a general effect of stress on heterosis linked to a stronger performance decline under warmer conditions than at sites of origin of populations. Our study therefore suggests that a general heterozygote deficit is more detrimental under temperature stress, and that interpopulation outcrossing can alleviate this effect.

A main result of this study is that expressed mutational load estimated by heterosis was not significantly enhanced under thermal stress. Indeed, while analyses on pot-level performance were supportive of an effect of the cross type, genomic load, and environmental stress on performance, these analyses did not support a three-way interaction between effects. This suggests that environmental stressfulness does not enhance the effect of the higher accumulation of recessive deleterious mutations on the expression of mutational load. Analyses on population-level heterosis were not clearly supportive of an interaction either. Even though the model including interactions between stress and genomic load was better supported than the model without interactions, the interaction was not significant in the *χ*^2^ test. This suggest that some variation in heterosis may be the result of an interaction, but other variation may easily blur the effect. Such environment dependence has often been evoked due to the similarities in the underlying genetics between heterosis (if caused by dominance) and inbreeding depression (Fenster & Galloway, 2000; Oakley et al., 2015). Indeed, a previous meta-analysis on the stress dependence of heterosis found that the benefit of outcrossing was considerably higher under stress (Frankham, 2015); however, the study only focused on small and inbred populations and did not include larger populations for comparison. Some of them may also show a heterotic effect under outbreeding that is enhanced under stress, as found in our study.

We think that our result for *A. lyrata* is robust despite caveats that were previously mentioned for the study of environment dependence of load. It was argued that environment dependence of the expression of load depends on the strength of stress (Fox & Reed, 2010). In our study, exposure to warmer conditions strongly reduced WPC population performance, by up to 92%. Type and novelty of stress were also suggested to affect stress dependence of the expression of load (Willi et al., 2007a; Sandner & Matthies, 2016). If *A. lyrata* had regularly experienced conditions we considered stressful here across its evolutionary history, deleterious alleles expressed under these conditions may have already been purged (Hedrick, 1994; Bijlsma et al., 1999; Enders & Nunney, 2016). However, the expression of mutational load was also not enhanced by stress based on relative performance, a broader measure of stress likely depicting also novel stressors.

Furthermore, the interaction between genomic load and stress could have been masked by outbreeding depression, previously reported to counteract heterosis under environmental stress such as drought (Prill et al., 2014). Outbreeding depression is expected to be stronger in crosses involving populations of highest divergence (Lynch, 1991; Fenster & Galloway, 2000; Oakley et al., 2015), although more recent studies find this relationship to be weak or absent in large datasets (Vasseur et al., 2019; Clo et al., 2021). In our study, partner populations came from the centres of geographic distribution and were unlikely highly divergent, given the shared post-glacial history and moderate climatic differences between populations. Furthermore, outbreeding depression was only found in few populations, mostly of the core of distribution, and it seemed unaffected by stress. The high performance gain, especially in populations with high load, also speaks in favour of an overwhelming positive effect of heterosis and little action of outbreeding depression. For these reasons, we conclude that the absence of significant environment dependence of mutational load seems real.

Heterosis was nonetheless affected by stress in our study, especially warm stress. Heterosis increased under warmer conditions than at the sites of origin of populations (Fig. 2). Perrier et al. (2020) previously linked patterns of heightened heterosis with increasing genomic load to a performance decline in WPC, with BPC being fairly constant across that gradient. Here we observed that WPC and BPC performance generally declined with increasing warm stress, but less in BPC, as suggested by the pot-level analysis on performance. Lower sensitivity to stress in outbred plants could result from recessive deleterious mutations being frequently heterozygous and their better buffering under stress by *e.g.* heat shock proteins (Rutherford & Lindquist, 1998; Queitsch et al., 2002; Bergman & Siegal, 2003). Higher performance of heterotic hybrids under stress in *A. thaliana* has been linked to disruptions of stress response pathways in inbred parental lines (Korn et al., 2010; Yang et al., 2015; Miller et al., 2015). Previous research in the context of agriculture or genetic rescue has also provided many examples of the effect of heterosis increased under stress (Frankham, 2015; Blum, 2013; Fujimoto et al., 2018). Here, the overall increase in heterosis under warm stress independent of genomic load suggests that most populations of *A. lyrata* suffer from a general heterozygote deficit whose expression is heat-sensitive.

Our results offer some important insights to the understanding of heterosis and its stress-dependence. First, model selection showed that the effect of increasing stressfulness was not linear but better described by a positive accelerating relationship. This was the case both when stress was expressed as the difference in temperature regime during a critical part of the year for *A. lyrata* between common garden and site of origin of populations or as the performance decline in a garden by populations. Second, when stress was depicted by temperature difference, it was the unsigned temperature difference that depicted stress better. While cooler conditions led to relatively high plant performance, it was the warmer conditions that were stressful and that led to higher heterosis. Third, even though interactions were not significant between genomic load and stressfulness on heterosis, our study suggests that they may still be of some relevance. One indication is that model selection pointed to the inclusion of interactions being a better model predicting heterosis. Another indication is that pot-level performance revealed significant negative interactions between genomic load and the linear term of stress, suggesting that populations with high genomic load performed worse under increasingly warmer conditions than at the site of origin, independent of cross type.

Our study has two main implications beyond the study of heterosis. The first is the role of expressed mutational load for range limits. Populations expanding beyond range limits may suffer from a twofold genetic Allee effect (Luque et al., 2016). A first is linked to the process of range expansion itself: in the absence of strong environmental gradients, range expansion may lead to the accumulation of high mutational load in populations at range limits (Peischl et al., 2013; Peischl et al., 2015; Gilbert et al., 2017). Further range expansion originating from these range-edge populations may therefore lead to an even higher accumulation of mutational load beyond tolerable levels to maintain population demographic rates (Perrier et al., 2020). A second genetic Allee effect is due to a general heterozygote deficit expressed under stressful, particularly warm-stress conditions. However, this problem was not confirmed for range-edge populations at range limits. Northern and southern range-edge populations did not significantly differ in heterosis when transplanted at their respective range edge or beyond. Nonetheless, the stress-dependent genetic Allee effect could be triggered by extreme environmental events that may be more frequent beyond range limits and by long-range dispersal into less suitable habitats than those immediately beyond range limits. Future simulation studies on range dynamics (Gilbert et al., 2017; Polechová, 2018) should therefore consider the consequences of stress-dependent genetic Allee effects on range expansion.

The second implication of our study is that interpopulation outbreeding can have a generally positive effect across populations of the distribution of a species. A great majority of populations indeed benefitted from outbreeding, especially under warmer conditions than what the populations had experienced in the past. Such heterotic effects can often be beneficial beyond the first generation (Willi et al., 2007b; Frankham, 2016). Our results therefore motivate the study of assisted gene flow in the context of climate warming to preserve species across the entire range. Such management may be worth being tested particularly for weak dispersers, potentially combined with the use of climate-adjusted provenances such as explored in restoration ecology (Prober et al., 2015). Populations at the southern range limits, with high genomic load, may also need to be supplemented with material from similar climatic conditions but different demographic history to maximize heterozygote advantage and ensure the local persistence of species.

## Acknowledgements

This work was supported by the Swiss National Science Foundation (31003 A_166322). We are grateful to Celia Evans (Paul Smith’s College, Paul Smith, NY), Joan Edwards (Williams College, Williamstown, MA), Heather Peckham Griscom (James Madison University, Harrisonburg, VA), William K. Smith (Wake Forest University, Winston-Salem, NC) and Rodney Mauricio (University of Georgia, Athens, GA) for logistical support in the US. For field assistance, we thank Mary Anderson, Michael Boyd, Bennet Coe, Scott Cory, Rachel Hillyer, Andrew Jones, Deidre Keating, Larry Kummer, David Lampman, Anastasia Levie-Sprick, Blake Macko, Shannon Malisson, Kathryn McGee, Althea Neighbors, Debra Rogers-Gillig, Caleb Rose, Amber Scarabaggio, Anna Shutley, Caroline Vath and Audrey Werner. For assistance with seed counts, we thank Olivier Bachmann, Markus Funk, and Susanna Riedl. Collection permits were provided by the Clinton County Conservation Board, Cornell University, Fort Leonard Wood Army Base, Iowa Department of Natural Resources, Missouri Department of Conservation, New York State Office of Parks, Ontario Parks, Palisades Interstate Park Commission, Rock Island Lodge, United States National Park Service, Virginia Department of Conservation and Recreation and the Wisconsin Department of Natural Resources.

## Author Contributions

All authors contributed to the study design. YW collected the seeds in the field, AP and DS raised and crossed plants. AP analyzed the data and wrote the manuscript, with support by YW.

## Data accessibility statement

All data is stored in Dryad (https://doi.org/10.5061/dryad.cc2fqz642).

