## Supplementary for "Environment-dependence of the expression of mutational load and species’ range limits"

**Supplementary Material**

**Table S1: Information on the *Arabidopsis lyrata* populations studied**

| Population | Latitude [° N] | Longitude [° W] | Ecological variable | Variables of population history | |
|---|---|---|---|---|---|
| | | | Min. temp. early spring [°C] † | Cluster | Genomic load ($P_n f_n / P_s f_s$) ‡ |
| IA1 | 41.97 | 90.37 | -0.9 | West | 0.80690 |
| IN1 | 41.61 | 87.19 | 0.0 | West | 0.83283 |
| MD2 | 38.99 | 77.25 | 3.8 | East | 0.78476 |
| MO1 | 37.72 | 92.06 | 4.5 | West | 0.90542 |
| MO2 | 38.47 | 90.71 | 4.0 | West | 1.03637 |
| NC2 | 36.04 | 81.16 | 3.8 | East | 0.91391 |
| NC4 | 36.41 | 79.96 | 5.0 | East | 0.86410 |
| NY1 | 41.3 | 73.98 | 0.4 | East | 0.77297 |
| NY4 | 42.35 | 76.39 | -2.5 | East | 0.77737 |
| NY5 | 42.66 | 74.02 | -3.0 | East | 0.78503 |
| NY6 | 43.00 | 76.09 | -2.1 | East | 0.77053 |
| ON1 | 42.87 | 79.18 | -1.7 | West | 0.96393 |
| ON11 | 48.77 | 87.13 | -7.9 | West | 1.09270 |
| ON12 | 49.65 | 94.92 | -7.8 | West | 0.85352 |
| ON3 | 43.26 | 81.84 | -2.6 | West | 0.86054 |
| ON8 | 47.93 | 84.85 | -7.5 | West | 0.88166 |
| PA3 | 41.28 | 77.87 | -1.4 | East | 0.76180 |
| VA1 | 37.42 | 77.02 | 5.5 | East | 0.81838 |
| WI1 | 43.83 | 89.72 | -3.3 | West | 0.73834 |
| WV1 | 38.96 | 79.29 | 1.1 | East | 0.82191 |

† Data extracted from WorldClim database version 2.0 (Fick & Hijmans, 2017). ‡ (Willi et al., 2018).

**Table S2: Summary of the crossing experiment and of the seeds sown in each common garden**

| Mother population | Father population | No. of cross families | No. of seeds sown in each common garden | | | | | Cross type |
|---|---|---|---|---|---|---|---|---|
| | | | CG1 | CG2 | CG3 | CG4 | CG5 | |
| NY1 | NY1 | 9 | 70 | 69 | 63 | 69 | 66 | WPC |
| NY5 | NY5 | 12 | 72 | 72 | 72 | 72 | 72 | WPC |
| IN1 | IN1 | 11 | 72 | 72 | 72 | 72 | 72 | WPC |
| MO1 | MO1 | 12 | 72 | 72 | 69 | 72 | 72 | WPC |
| ON11 | ON11 | 12 | 72 | 69 | 66 | 72 | 66 | WPC |
| ON12 | ON12 | 12 | 69 | 69 | 72 | 72 | 72 | WPC |
| MO2 | MO2 | 12 | 72 | 72 | 72 | 72 | 72 | WPC |
| NC2 | NC2 | 12 | 69 | 70 | 72 | 72 | 72 | WPC |
| NC4 | NC4 | 11 | 72 | 72 | 72 | 72 | 72 | WPC |
| IA1 | IA1 | 9 | 70 | 72 | 57 | 72 | 66 | WPC |
| VA1 | VA1 | 9 | 72 | 72 | 72 | 72 | 72 | WPC |
| MD2 | MD2 | 10 | 69 | 69 | 69 | 72 | 69 | WPC |
| WV1 | WV1 | 12 | 69 | 69 | 66 | 72 | 60 | WPC |
| PA3 | PA3 | 11 | 72 | 72 | 72 | 72 | 72 | WPC |
| NY6 | NY6 | 12 | 72 | 72 | 66 | 72 | 63 | WPC |
| NY4 | NY4 | 10 | 70 | 69 | 69 | 72 | 72 | WPC |
| WI1 | WI1 | 10 | 66 | 72 | 72 | 72 | 72 | WPC |
| ON8 | ON8 | 6 | 27 | 30 | 27 | 37 | 27 | WPC |
| ON3 | ON3 | 8 | 69 | 69 | 69 | 72 | 66 | WPC |
| ON1 | ON1 | 12 | 72 | 72 | 72 | 72 | 69 | WPC |
| NY5 | NY1 | 12 | 72 | 72 | 72 | 72 | 72 | BPC |
| IN1 | IA1 | 11 | 72 | 69 | 69 | 72 | 72 | BPC |
| MO1 | IA1 | 12 | 72 | 72 | 72 | 72 | 72 | BPC |
| ON11 | IA1 | 9 | 55 | 54 | 54 | *0* | 72 | BPC † |
| ON12 | IA1 | 10 | 72 | 72 | 72 | 72 | 72 | BPC |
| MO2 | IA1 | 12 | 72 | 72 | 72 | 72 | 72 | BPC |
| NC2 | NY1 | 12 | 69 | 70 | 69 | 69 | 72 | BPC |
| NC4 | NY1 | 10 | 72 | 72 | 72 | 72 | 72 | BPC |
| VA1 | NY1 | 10 | 72 | 72 | 72 | 78 | 72 | BPC |
| MD2 | NY1 | 9 | 72 | 72 | 72 | 72 | 72 | BPC |
| WV1 | NY1 | 11 | 72 | 72 | 66 | 72 | 66 | BPC |
| PA3 | NY1 | 11 | 72 | 72 | 72 | 72 | 72 | BPC |
| NY6 | NY1 | 11 | 72 | 72 | 72 | 72 | 72 | BPC |
| NY4 | NY1 | 11 | 72 | 72 | 72 | 72 | 72 | BPC |
| WI1 | IA1 | 11 | 78 | 72 | 72 | 72 | 72 | BPC |
| ON8 | IA1 | 5 | 34 | 35 | 33 | 39 | 33 | BPC |
| ON3 | IA1 | 10 | 63 | 63 | 63 | 72 | 63 | BPC |
| ON1 | IA1 | 12 | 72 | 72 | 72 | 72 | 69 | BPC |

Total number of seeds sown: 12,933; total number of cross-combinations: 401; total number of population-level WPC: 100; total number of population-level BPC: 89. † Offspring between ON11 and IA1 are missing in CG4 due to too low numbers of seeds. Heterosis was not calculated for this cross-site-combination.

**Table S3: Information on the common garden sites**

| Transplant site | Location | Latitude [° N] | Longitude [° W] | Min. temperatures early spring (°C) † |
|---|---|---|---|---|
| CG1 (NY) | Beyond northern edge | 44.51 | 74.02 | -6.5 |
| CG2 (MA) | Northern edge | 42.72 | 73.22 | -3.3 |
| CG3 (VA) | Center | 38.43 | 78.86 | 1.8 |
| CG4 (NC) | Southern edge | 36.13 | 80.28 | 5.6 |
| CG5 (GA) | Beyond southern edge | 33.93 | 83.36 | 8.0 |
| Mean northern pop. ‡ | North | - | - | -2.5 |
| Mean center pop. ‡ | Center | - | - | 1.1 |
| Mean southern pop. ‡ | South | - | - | 4.7 |

† Data extracted from WorldClim database version 2.0 (Fick & Hijmans, 2017); ‡ Data measured for mean population of the eastern cluster. Northern populations: NY4, NY5, NY6; center populations: PA3, MD2, WV1; southern populations: VA1, NC2, NC4.

**Table S4: Average minimum temperature in early spring (March and April) at the garden sites ($T_{min\ CG}$)**

| Site | $T_{min}$ year 2 [° C] | $T_{min}$ year 2 + 3 [° C] |
|---|---|---|
| CG1 (NY) | -5.8 | -6.0 |
| CG2 (MA) | -2.8 | -1.7 |
| CG3 (VA) | 2.2 | † |
| CG4 (NC) | 5.2 | 7.0 |
| CG5 (GA) | 6.4 | 7.0 |

Temperature recordings from data loggers (iButton®, Maxim Integrated Products, Inc) placed in each common garden site (1.5 m above ground, in the shadow). † In the analysis, replaced by the value of year 2.

**Table S5: Summary of population-level heterosis or performance of within- (WPC) and between-population crosses (BPC) based on *multiplicative performance* (*MP*), or the finite rate of increase (*λ*), at the five common garden sites**

| | Population heterosis | | | | Population performance | | | | | |
|---|---|---|---|---|---|---|---|---|---|---|
| | | | | | $WPC_{individual\ population}$ | | $WPC_{mean\ population}$ | | BPC | |
| **Dependent variable** | ***N*** | ***Min.*** | ***Mean*** | ***Max*** | ***N*** | ***Mean*** | ***N*** | ***Mean*** | ***N*** | ***Mean*** |
| MP | 89 | -0.96 | 1.88 | 23.50 | 89 | 40.36 | 100 | 41.31 | 89 | 75.68 |
| *λ* | 89 | -0.53 | 0.73 | 7.29 | 89 | 0.98 | 100 | 1.00 | 89 | 1.38 |

Performance of WPC was calculated individualy for each partner or target population ($WPC_{individual\ population}$), or as average between each target and partner population ($WPC_{mean\ population}$).

**Table S6: Model selection on potential predictors of heterosis (models sorted by ascending AICc values)**

| **Model** | ***AICc*** | ***Δi*** | ***wi*** |
|---|---|---|---|
| Genomic load + stress + stress$^2$ + (genomic load * stress) + (genomic load * stress$^2$ ) | 112.9 | 0.0 | 0.96 |
| Genomic load + stress + stress$^2$ | 119.6 | 6.7 | 0.03 |
| Genomic load | 122.6 | 9.7 | 0.01 |
| Genomic load + stress | 128.1 | 15.2 | 0.00 |
| Genomic load + \|stress\| | 130.2 | 17.3 | 0.00 |
| Genomic load + stress + (genomic load * stress) | 132.6 | 19.7 | 0.00 |
| Genomic load + \|stress\| + (genomic load * \|stress\|) | 133.8 | 20.9 | 0.00 |

The dependent variable was heterosis based on *multiplicative performance*. Genomic load was the ratio of genome-wide non-synonymous to synonymous polymorphic sites adjusted by their mean derived frequency (Willi et al., 2018), and stress was the difference in minimum temperature in early spring between common garden and site of origin ($\Delta\ T_{\text{min}} = T_{\text{min CG}} - T_{\text{min origin}}$). Models are sorted by ascending AICc values, with lower *AICc* values indicating a better fit. The difference between the best fit model and the others is indicated as *Δi*, while the weight of each model is indicated by *wi*.

**Table S7: Results of model testing for the effect of the genomic estimate of mutational load, environmental stress ($\Delta T_{min} = T_{min\ CG} - T_{min\ origin}$) and their interaction on *multiplicative performance* (*MP*) of within-population crosses (WPC), at the five common garden sites**

| | | Genomic load (GL) | | Stress (S) | | Stress$^2$ ($S^2$) | | GL * S | | GL * $S^2$ | | | |
|---|---|---|---|---|---|---|---|---|---|---|---|---|---|
| **Dependent variable** | ***N*** | ***β*** | ***$\chi^2$*** | ***β*** | ***$\chi^2$*** | ***β*** | ***$\chi^2$*** | ***β*** | ***$\chi^2$*** | ***β*** | ***$\chi^2$*** | ***$R^2m$*** | ***$R^2c$*** |
| MP | 89 | **-1.42** | **6.99 *** | **-2.64** | **9.76 **** | **-1.26** | **11.22 ***** | -5.42 | 2.29 | 6.38 | 3.61 (*) | 0.38 | 0.72 † |

Performance estimates were an average over within-population crosses of pairs of target and partner populations in a common garden (based on within-population averages). Values were $\log_{10}$-transformed prior to analysis. Stress, $\Delta T_{min}$, was an average over target and partner population. Each model was optimized with the *bobyqa* optimizer to improve convergence. Test statistics include regression coefficient ($\beta$), $\chi^2$-value of each fixed effect, and the marginal and conditional $R^2$ of the model. Model fits with significant (positive) intercept are indicated by †. Regression coefficients with $P$-values < 0.05 are written in bold; significance is indicated: (*) $P < 0.1$, * $P < 0.05$, ** $P < 0.01$, *** $P < 0.001$. Results for random effects are not shown. For one of the five common gardens (CG3), the experiment stopped early and the dependent variable considers performance to year 2 only.

**Table S8: Magnitude of effect of the genomic estimate of mutational load and environmental stress (Δ $T_{min}$ = $T_{min\ CG}$ - $T_{min\ origin}$) on ($log_{10}$-transformed) *multiplicative performance (*MP) of within-population crosses (WPC) (in %), at the five common garden sites**

| | | | Stress | |
|---|---|---|---|---|
| **Dependent variable** | ***N*** | ***Genomic load*** | ***Warmer CG*** | ***Colder CG*** |
| MP | 89 | -74.8 (0.74; 1.09) | -92.0 (0; 13.5) | 10,8 (0; -8.8) |

For the general model ofTable S7 analyzing WPC performance, the magnitude of effect of genomic load and environmental stress was calculated as: percentage difference between the back-transformed predicted performance corresponding to the maximal value of the predictor variable (in parenthesis, right) in our sampling and the back-transformed predicted performance corresponding to the minimal value of the predictor variable (in parenthesis, left). Stress, Δ $T_{min}$ was an average over target and partner population. The magnitude of the effect of genomic load was estimated considering temperatures close to those of the site of origin (stress = 0). The magnitude of the effect of environmental stress was estimated considering mean genomic load values and calculated separately if the common gardens (CG) were warmer or colder than the site of origin of the population.

**Table S9: Model selection on potential predictors of heterosis (models sorted by ascending AICc values)**

| **Model** | ***AICc*** | ***Δi*** | ***wi*** |
|---|---|---|---|
| Genomic load + stress + $stress^2$ + (genomic load * stress) + (genomic load * $stress^2$) | 111.9 | 0.0 | 0.96 |
| Genomic load + stress + $stress^2$ | 118.4 | 7.3 | 0.04 |
| Genomic load | 122.6 | 11.5 | 0.00 |
| Genomic load + stress + (genomic load * stress) | 124.7 | 13.6 | 0.00 |
| Genomic load + stress | 125.6 | 14.6 | 0.00 |

The dependent variable was heterosis based on *multiplicative performance I*. Genomic load was the ratio of genome-wide non-synonymous to synonymous polymorphic sites adjusted by their mean derived frequency (Willi et al., 2018), and stress was the relative difference of within-population cross performance of each population at a site compared to the best site (1 - $W_{\mathrm{WPC}}/W_{\mathrm{max}}$). Models are sorted by ascending AICc values, with lower *AICc* values indicating a better fit. The difference between the best fit model and the others is indicated as *Δi*, while the weight of each model is indicated by *wi***.**

**Table S10: Result of model testing for the effect of the genomic estimate of mutational load, stress based on relative performance of within-population crosses (1 - $W_{WPC}/W_{max}$), and their interaction on heterosis based on *multiplicative performance* (MP), or the finite rate of increase ($\lambda$), at the five common garden sites**

| | | Genomic load (GL) | | Stress (S) | | Stress² (S²) | | GL * S | | GL * S² | | | |
|---|---|---|---|---|---|---|---|---|---|---|---|---|---|
| **Dependent variable** | ***N*** | ***β*** | ***χ²*** | ***β*** | ***χ²*** | ***β*** | ***χ²*** | ***β*** | ***χ²*** | ***β*** | ***χ²*** | ***R²m*** | ***R²c*** |
| MP | 89 | **1.92** | **6.66 **** | 0.64 | 1.59 | **1.21** | **6.88 **** | -1.47 | 0.09 | -1.04 | 0.03 | 0.18 | 0.44 †, ‡ |
| $\lambda$ | 89 | 0.67 | 3.02 (*) | **0.92** | **14.20 ***** | **0.48** | **3.97 *** | 1.04 | 0.33 | 3.15 | 2.97 (*) | 0.29 | 0.60 † |

Estimates of population heterosis were $\log_{10}$-transformed prior to analysis. Each model was optimized with the *bobyqa* optimizer to improve convergence. Test statistics include regression coefficient ($\beta$), $\chi^2$-value of each fixed effect the marginal and conditional $R^2$ of the model. Model fits with significant (positive) intercept are indicated by †. Regression coefficient with *P*-values < 0.05 are written in bold; significance is indicated: : (*) $P < 0.1$, * $P < 0.05$, ** $P < 0.01$, *** $P < 0.001$. For one of the five common gardens (CG3), the experiment stopped early and variables consider performance to year 2 only (‡).

**Table S11: Result of model testing for the effect of transplant site relative to the species' distribution (beyond the edge compared to edge [0]) within each edge type (northern or southern), on heterosis based on *multiplicative performance* (*MP*), and the finite rate of increase (*λ*), at the five common garden sites**

| | **Northern edge** | | | | | **Southern edge** | | | | |
|---|---|---|---|---|---|---|---|---|---|---|
| **Dependent variable** | ***N*** | ***β*** | ***χ²*** | ***R²m*** | ***R²c*** | ***N*** | ***β*** | ***χ²*** | ***R²m*** | ***R²c*** |
| MP | 12 | -0.10 | 0.33 | 0.03 | 0.03 | 12 | -0.05 | 0.03 | 0.00 | 0.00 † |
| λ | 12 | -0.05 | 0.55 | 0.05 | 0.05 | 12 | 0.01 | 0.00 | 0.00 | 0.00 |

Estimates of population heterosis were $\log_{10}$-transformed prior to analysis. Northern and southern edge were analyzed in separate models. Each model was optimized with the *bobyqa* optimizer to improve convergence. Test statistics include regression coefficient ($\beta$), $\chi^2$-value of the effect of the transplant site and the marginal and conditional $R^2$ of the model. Model fits with significant (positive) intercept are indicated by †. Regression coefficient with $P$-values < 0.05 are written in bold; significance is indicated: (*) $P < 0.1$, * $P < 0.05$, ** $P < 0.01$, *** $P < 0.001$. Results for random effects are not shown.

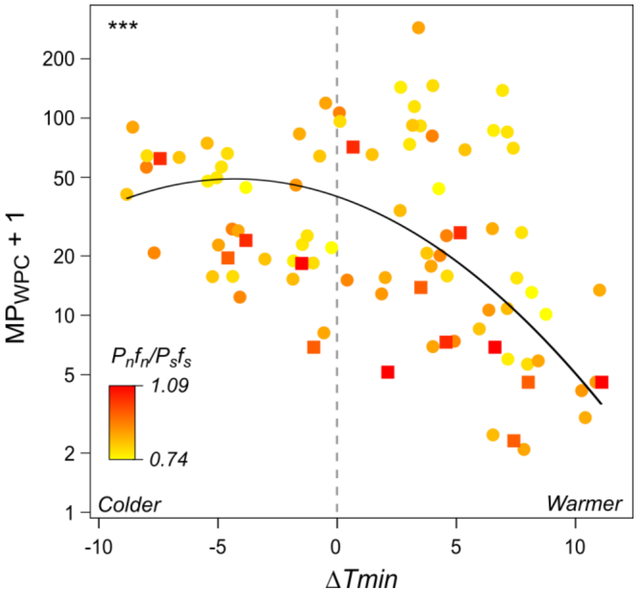


**Figure S1: Stress dependence within-population cross (WPC) performance.** Performance was based on multiplicative performance (MP) of within-population crosses averaged over pairs of target and partner populations in a common garden. Outcrossing populations are indicated by dots, selfing populations by squares. Stress was depicted as the difference in minimum temperature in early spring between common garden and the site of origin of a population ($\Delta\ T_{\mathrm{min}} = T_{\mathrm{min\ CG}} - T_{\mathrm{min\ origin}}$; positive values to the right of the vertical dashed line indicate a warmer environment), averaged over pairs of target and partner populations. The black lines represent model-predicted relationships between performance and environmental stress (test statistics inTable S7; *** $P < 0.001$). Genomic mutational load, the ratio of genome-wide non-synonymous to synonymous polymorphic sites adjusted by their mean derived frequency, $P_n f_n/P_s f_s$ (Willi et al., 2018), is represented in shades of yellow (low) to red (high).

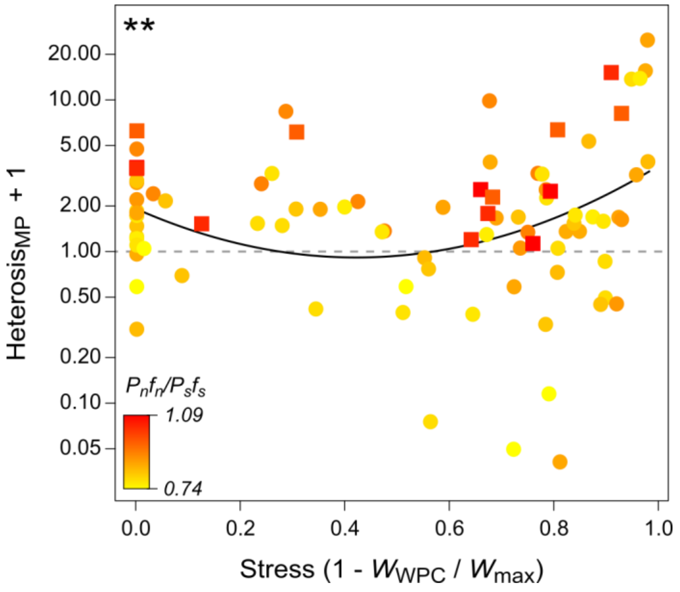


**Figure S2: Stress dependence of the expression of mutational load estimated by heterosis.** Population heterosis was estimated based on multiplicative performance (MP). Outcrossing populations are indicated by dots, selfing populations by squares. Stress was the relative decline in performance of within-population crosses in a garden relative to the garden of highest performance for that population ($1 - W_{WPC}/W_{max}$). The horizontal dashed lines indicates when heterosis drops below 0 and outbreeding depression dominates. The black line represents the model-predicted slope for heterosis (test statistics inTable S10, ** $P < 0.01$). Genomic mutational load, the ratio of genome-wide non-synonymous to synonymous polymorphic sites adjusted by their mean derived frequency, $P_n f_n / P_s f_s$ (Willi et al., 2018), is represented in shades of yellow (low) to red (high).

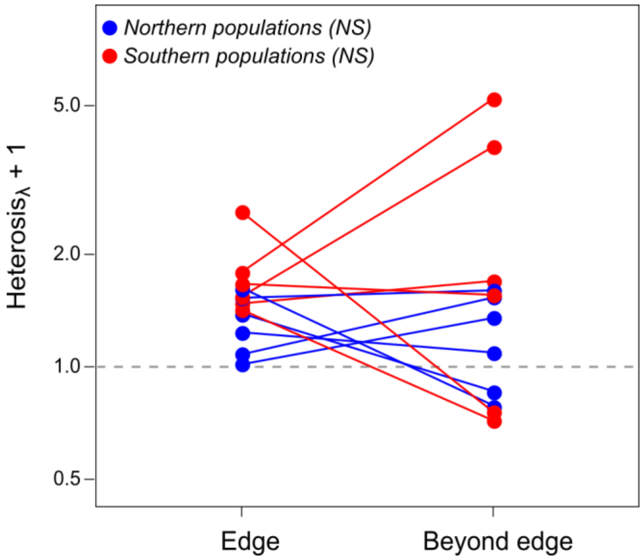


**Figure S3: Expression of mutational load at range limits and beyond.** Population heterosis was estimated based on the finite rate of increase, $\lambda$. The horizontal dashed lines indicates when heterosis drops below 0 and outbreeding depression dominates. The effect of transplanting beyond the range edge was tested for both northern (blue) and southern (red) populations at their respective northern and southern edge (CG2, CG5) and beyond the edge (CG1, CG5). Lines indicate the change in heterosis of each population. Northern and southern edge were analyzed in separate models. Test statistics are reported in Table S11. No significant effect of transplanting beyond the range edge was found, neither for northern nor for southern populations (NS).

### Methods S1: Parametrization of priors, and hierarchical mixed-effects model analyzed in a Bayesian (MCMC) framework, with individual *multiplicative performance I* as dependent variable

#### S1A: Priors

Priors were set to be weak, using parameter expansion to improve convergence. *R* specifies the priors for the fixed effects, *G* specifies the priors for the random effects.

```
priors.model=list(
  R=list(V=diag(2), n=1, fix = 2),
  G=list(G1=list(V=diag(2), n=2, alpha.mu = rep(0,2),alpha.V = diag(2)*25^2),
        G2=list(V=diag(4), n=4, alpha.mu = rep(0,4),alpha.V = diag(4)*25^2),
        G3=list(V=diag(2), n=2, alpha.mu = rep(0,2),alpha.V = diag(2)*25^2),
        G4=list(V=diag(4), n=4, alpha.mu = rep(0,4),alpha.V = diag(4)*25^2),
        G5=list(V=diag(2), n=2, alpha.mu = rep(0,2),alpha.V = diag(2)*25^2),
        G6=list(V=diag(2), n=2, alpha.mu = rep(0,2),alpha.V = diag(2)*25^2)))
```

#### S1B: Parametrization of hierarchical mixed-effects models analyzed in a Bayesian framework

Multiplicative performance was split into two parts: the *zero_part*, a binary transformation of performance with *zero_part* = 1 if performance > 0, or else *zero_part* = 0; and the *norm_part* containing only the $\log_{10}$ transformed performance measures if *zero_part* = 1.

```
model = MCMCglmm(cbind(norm_part, zero_part) ~ trait -1 + trait:cross type * trait:genomic load
* trait:poly(stress, degree = 2),
            random = ~ us(trait):maternal population
            + us(trait:cross type):maternal population
            + us(trait): maternal population: maternal family
            + us(trait: cross type):maternal population:maternal family
            + us(trait):common garden + us(trait):common garden:block,
            prior = priors.model, rcov = ~idh(trait):units,
            family=c('gaussian', 'categorical'),
            burnin = 5000, thin = 100, nitt = 50000, data=data)
```

Convergence was assessed using Gelman-Rubin diagnostics and Gelman-Rubin-Brooks plots with the functions *gelman.diag* and *gelman.plot* implemented in the *MCMCglmm R* package (Hadfield, 2010; Hadfield, 2019), and autocorrelation was assessed by estimating non-independence between successive samples for means and variances with the function *autocorr* in *MCMCglmm*.

**Methods S2: Parametrization of the hierarchical mixed-effects models, with heterosis as dependent variable**

**S2A: Hierarchical mixed-effects model testing for the effect of stress and genomic load**

```
Model = lmer(log10(heterosis + 1) ~ genomic load * poly(stress, degree = 2)
        + (1 | maternal population)
        + (1 | common garden),
        data = data)
```

**S2B: Hierarchical mixed-effects model testing for the effect of transplant**

```
Model = lmer(log10(heterosis + 1) ~ transplant site
        + (1 | maternal population),
        data = data)
```

**Appendix S1:**

Two additional factors could affect heterosis, namely genetic incompatibility likely reflected by neutral genetic divergence, and environmental divergence due to divergent adaptation between target and partner populations. Their effect was explored as additional fixed effects in initial model selection analyses of the model described in Method S6A. Genetic divergence was estimated by performing a PCA on a SNP dataset generated from a previously published dataset of whole-genome sequence data (Willi *et al.*, 2018). Environmental divergence was estimated as the difference in average minimum temperature in March and April between target and partner populations based on the WorldClim 2.0 database (Fick & Hijmans, 2017). Genetic divergence was correlated with genomic load ($r$ = 0.57, $N$ = 18) and environmental divergence was correlated with environmental stress ($r$ = 0.39, $N$ = 89). Neither genetic divergence nor environmental divergence were significant both in models where they were considered directly and when their residuals were used after accounting for genomic load and environmental stress, respectively, and AICcs were higher (data not shown).
